# Gaussian Mixture Modeling Extensions for Improved False Discovery Rate Estimation in GC-MS Metabolomics

**DOI:** 10.1101/2023.02.06.527348

**Authors:** Javier E. Flores, Lisa M. Bramer, David J. Degnan, Vanessa L. Paurus, Yuri E. Corilo, Chaevien S. Clendinen

## Abstract

The ability to reliably identify small molecules (e.g. metabolites) is key towards driving scientific advancement in metabolomics. Gas chromatography–mass spectrometry (GC-MS) is an analytic method that may be applied to facilitate this process. The typical GC-MS identification workflow involves quantifying the similarity of an observed sample spectrum and other features (e.g. retention index) to that of several references, noting the compound of the best-matching reference spectrum as the identified metabolite. While a deluge of similarity metrics exists, none quantify the error rate of generated identifications, thereby presenting an unknown risk of false identification or discovery. To quantify this unknown risk, we propose a model-based framework for estimating the false discovery rate (FDR) among a set of identifications. Extending a traditional mixture modeling framework, our method incorporates both similarity score and experimental information in estimating the FDR. We apply these models to identification lists derived from across 548 samples of varying complexity and sample type (e.g. fungal species, standard mixtures, etc.), comparing their performance to that of the traditional Gaussian mixture model (GMM). Through simulation, we additionally assess the impact of reference library size on the accuracy of FDR estimates. In comparing the best performing model extensions to the GMM, our results indicate relative decreases in median absolute estimation error (MAE) ranging from 12% to 70%, based on comparisons of the median MAEs across all hit-lists. Results indicate that these relative performance improvements generally hold despite library size, however FDR estimation error typically worsens as the set of reference compounds diminishes.

For TOC graphic only

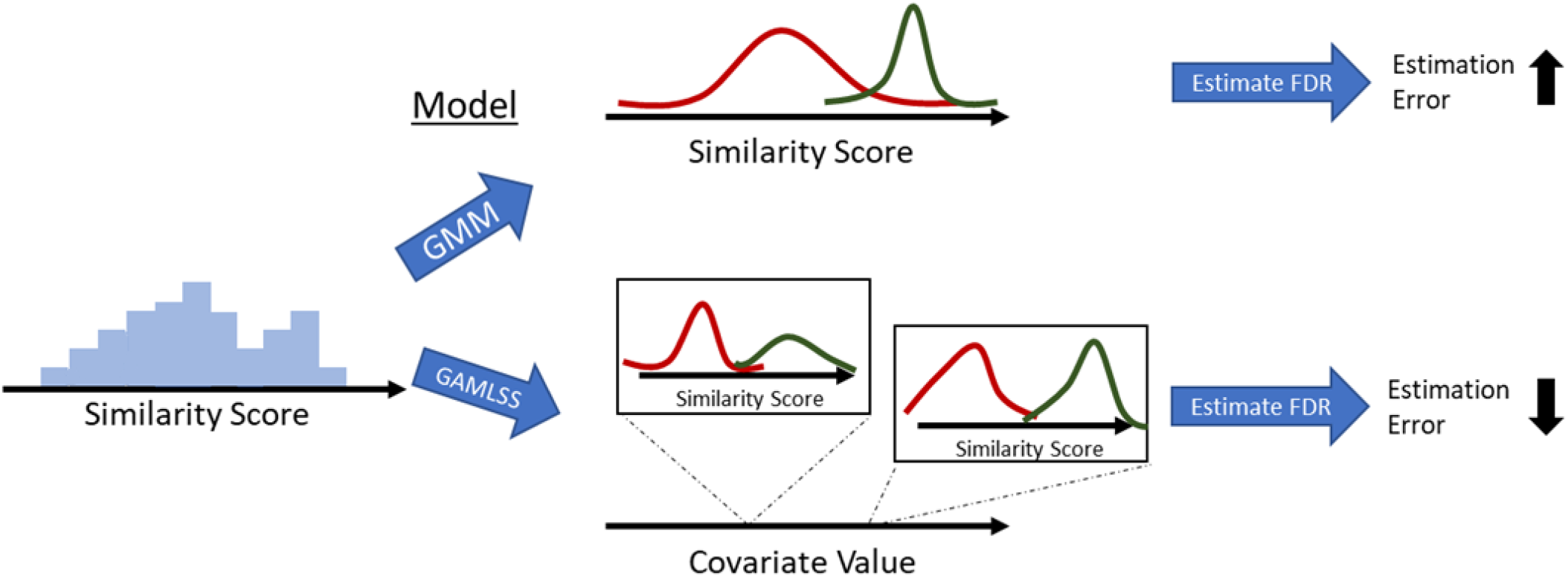

## 1. INTRODUCTION

Metabolites – which include the diverse set of hormones, signaling molecules and other small molecules – help paint a detailed picture of the chemical processes that govern biological systems. Thus metabolomics, the study of metabolites and other small molecules, impacts scientific advancement across a wide variety of biological and chemical research applications. Fundamental to these impactful metabolomic contributions, however, is the ability to accurately identify the set of metabolites in samples using a variety of analytical techniques.

Gas chromatography-mass spectrometry (GC-MS) is an analytic method that yields molecular fingerprints in the form of spectra, thus making GC-MS an ideal tool for metabolite identification. A single sample run through the GC-MS instrument generates numerous spectra to be identified. These observed or query spectra are then typically compared to those of known reference compounds, with each comparison being scored by a metric assessing their degree of similarity. The reference metabolite of each query spectrum’s best scoring match is then assigned as the spectrum’s identification, provided that the measured similarity is above the threshold level specified by the analyst. For example, when using the cosine dot-product, threshold levels between 0.6 and 0.7 are often used (1). Despite the use of these thresholds, there is no guarantee that the resulting set of identifications are without false positives and thus there is rarely an accurate estimate of the expected number of false identifications relative to the total set (i.e. false discovery rate, FDR).

Methods for false discovery rate estimation are commonly applied in proteomics and genomics, with some of these approaches utilizing biological structural information to inform or generate their estimates. In proteomics, for example, spectra are matched against a composite database of decoy and reference proteins, where decoys are generated by shuffling the reference protein sequences. The number of false positives is then estimated by doubling the number of matched decoys (2). Given the structural diversity of metabolites, such methods are not directly transferable. However, for liquid chromatography tandem mass spectrometry (LC-MS/MS), an alternative solution has been offered by Sheubert et al. who developed targetdecoy approaches with decoys generated through fragmentation trees (3–5). Yet the best performing of these approaches was found to only marginally outperform an empirical Bayes approach (6) when applied to noise-filtered data and substantially underperform otherwise.

The previous target-decoy-based approaches removed, existing FDR methods in metabolomics are largely probabilistic or model-based (1,7–9). However, the widespread adoption of these approaches has been hindered due to either their complexity, limitation(s) towards implementation, or their generalizability. As an example, the Basic Local Alignment Search Tool method requires that users perform several complex preprocessing steps and specify several parameters that influence resulting estimates (1). Kim and Zhang (9) offer a much less complicated solution based on similarity score differences, but their approach is based on a limited set of empirical trends observed when applying the weighted cosine correlation to the NIST 08 Mass Spectral Library data and has not been generalized to other scores or datasets.

Recognizing the need for an effective, relatively simple and generalizable approach to FDR estimation in GC-MS metabolomics, we propose models based on a conceptually intuitive Gaussian mixture (i.e. empirical Bayes) model (GMM, 6) with extensions that leverage both similarity score and experimental information towards FDR estimation. Furthermore, implementation of the proposed approach requires little to no pre-processing beyond that required by the standard GC-MS identification workflow. We compare the proposed models to the standard GMM across a total of 812 identification lists generated from 28 different similarity metrics applied separately to each of 29 different datasets of varying sample type and complexity. In addition, we conduct a simulation study to assess the potential impact of reference library size on the accuracy of FDR estimates generated by the considered approaches.

## 2. EXPERIMENTAL SECTION

### 2.1 Data

Our data were compiled from an internal Pacific Northwest National Laboratory (PNNL) repository of Agilent .D files collected over four years and processed by CoreMS (10), software developed at PNNL. Datafiles corresponding to standards samples were generated from standard metabolites purchased from Sigma Aldrich, and these standards were derivatized using a modified version of the FiehnLib protocol (11) as described in other works (12, 13). Briefly, dried samples underwent methoxyamination and trimethylsilyation (TMS) before being analyzed by an Agilent GC 7890A coupled with a single quadrupole MSD 5975C (Agilent Technologies). Data were collected over a mass range of 50-550 m/z, and a standard mixture of fatty acid methyl esters (FAMEs) (C8 – C28) were analyzed with samples to facilitate retention index alignment.

The processed and compiled data are representative of six categorizations of sample type of varying complexity: standards mixtures, human cerebrospinal fluid (CSF), human blood plasma, human urine, fungi species (*A. niger, A. nidulans, T. reesei*), and soil crust. Each of the human samples (i.e. CSF, plasma, and urine) contained deuterated internal standards (14). Within each category of sample type are collections of datasets, each of which are composed of spectra measured at different observed retention times/indices from across separate samples. In processing these data with CoreMS, a generous retention index window of +/- 35 was used to append lists of candidate metabolite matches to the corresponding processed spectra.

Candidate matches of each spectrum were then manually annotated as either “True Positive” (TP), “True Negative” (TN), or “Unknown” based on the assessment of expert analysts. Expert annotations were partly informed by a few guidelines. Specifically, for standards samples, any small compound that was not included in the mixture was labeled as a TN. Exceptions to this rule include FAMES and other commonly occurring compounds such as carbonate, phosphate, phosphoric acid, glycerol, and propylene glycol. In complex samples, for a spectrum with a candidate metabolite labeled as TP, all other candidate metabolites were labeled as TN. Lists of prevalent compounds within certain sample types were derived from the Human Metabolome Database (15) and CEU Mass Mediator (16). These lists were used to assign TN labels to candidate metabolites that were foreign to a particular sample type (e.g. benzene-1,2,4-triol in blood plasma), except for prevalent sugars, amino acids, nucleic acids, small organic compounds, and the FAMES and commonly occurring compounds previously mentioned. Candidate metabolites which could not be confidently assigned as either TP or TN were labeled as “Unknown”. Table 1 displays the number of datasets, samples, and total spectra corresponding to each sample type. Counts and percentages of all corresponding annotations (TP, TN, or Unknown) are also included. These datasets may be accessed at https://doi.org/10.25584/PNNL.data/1902325.

**Table 1.** Dataset, Sample, Spectrum, and Annotation Counts by Sample Type.

| Sample Type | # Datasets | # Samples | # Spectra | # (%) TP | # (%) TN | # (%) Unknown |
| --- | --- | --- | --- | --- | --- | --- |
| Standards | 11 | 140 | 10807 | 433 (0.09) | 452708 (96.19) | 17498 (3.72) |
| Human CSF | 8 | 206 | 25035 | 6437 (0.44) | 947132 (64.90) | 505832 (34.66) |
| Human Blood Plasma | 3 | 90 | 15621 | 2593 (0.32) | 333239 (41.46) | 467877 (58.22) |
| Human Urine | 2 | 66 | 22978 | 2340 (0.20) | 481163 (41.43) | 678008 (58.37) |
| Fungi | 3 | 30 | 8766 | 1102 (0.24) | 260721 (57.40) | 192407 (42.36) |
| Soil | 2 | 16 | 3622 | 581 (0.34) | 100457 (58.50) | 70688 (41.16) |

### 2.2 FDR Estimation

Estimates of the false discovery rate are generated through each of four general approaches. The first approach, a gaussian mixture model (GMM) serves as a baseline for comparison given its ubiquitous application as an FDR estimation method across a wide variety of scientific domains. The choice of GMM for baseline was also influenced by a separate simulation study indicating superior performance of the GMM as compared to the method introduced by Jeong et al (8). A description of this simulation study and its main finding are provided in the supplement (S1, Figure S1).

Within the context of GC-MS metabolomics, a GMM is fit to the measured similarity scores of the top-matched candidate metabolites from across a group of spectra. The fitted GMM presumes that the observed distribution of similarity scores arises from an unknown mixture of two latent populations, each normally distributed with some mean and variance. In the present context, these two latent populations correspond to the correctly matched metabolites (true positives) and incorrect matches (true negatives). The expectation-maximization algorithm (17) is often used to fit GMMs, and it provides an estimate of the unknown mixing proportion between the two latent populations and estimates of each population’s mean and standard deviation. These parameter estimates may then be used to assign each match a probability indicating their likelihood of inclusion among either of the two latent populations. Assuming larger similarity scores indicate a better-quality match, and given a set of *n* spectra with a similarity score threshold *t*, the FDR is estimated as

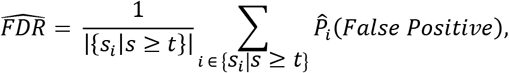

where *s_i_* = {*s*_1_, …, *s_n_*} are the measured similarity scores of the top matches for each of the *n* spectra, |{*s_i_*|*s* ≥ *t*}| is the number of top matches with scores greater than or equal to the threshold, and 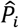 (*False Positive*) is the GMM estimated probability that the match of the *i^th^* spectrum belongs to the false positive population. If it is the case that smaller similarity scores indicate a better-quality match, the above computation remains the same, but each instance of *s ≥ t* is replaced with *s* ≤ *t*.

The remaining three modeling approaches that we consider are extensions to the GMM that are derived from a more general class of modeling framework first introduced by Rigby and Stasinopoulos (18). These extensions blend a regression-type framework with the GMM, effectively allowing one to model select GMM parameters (i.e. mixing probability, population means) as a linear function of one or more additional factors (i.e. covariates). The first extension, hereafter referred to as “Extension 1”, models the *mixing proportion* as a function of covariate(s), meaning that a match’s probability of belonging to a given latent population varies depending on the specified set of external factors. The second extension, hereafter referred to as “Extension 2”, models the underlying *means* of each latent population as a linear function of covariate(s), which implies that the degree of separation between the true and false positive populations changes according to the values of these factor(s). The final extension, hereafter referred to as “Extension 3”, is a combination of the first two, modeling the *mixing proportion* and *population means* through a linear function of covariates. Though each extension differs from the GMM in their construction and flexibility, each are like the GMM in that they ultimately yield probabilities of latent population membership for each match. Thus, FDR estimates are obtained through the same method as for the GMM. Figure 1 visually summarizes the conceptual differences between each of the four considered modeling approaches for FDR estimation.

**Figure 1.**
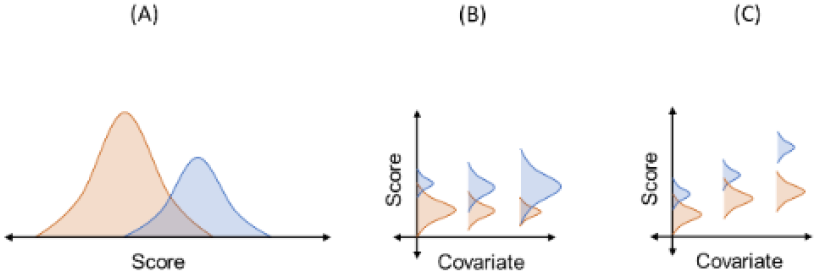
Conceptual differences between the standard GMM (A), first extension (B), and second extension (C). The third extension (not pictured) is a combination of (B) and (C). Changing mixing proportion are represented by an increase (decrease) in the relative sizes of each bell curve. Neither the mixing proportion or population means of the standard GMM (A) vary by covariate(s).

### 2.3. Model Covariates

Given that the proposed GMM extensions are characterized by the ability to model select parameter(s) by a set of factors, we consider all possible subsets of seven factors in the fitting of each model extension. These factors may be classified as either spectrum-specific or match-specific factors.

Spectrum-specific factors are those that vary across a sample query spectra. These include the number of candidate (i.e. potential) matches returned by CoreMS that were within the +/- 35 RI window (“PoMatchCnt”); the interference and peak height of the spectrum (“Intrfrnce”, “PeakHgt”); and the difference in similarity score between the top two ranked matches for the spectrum (“ScrDiff’). If only a single candidate match was returned by CoreMS, “ScrDiff’ was computed as either the difference between the measure score and 0 (when larger scores indicate better matches) or as the difference between the measured score and the largest empirically observed score across all spectra in the dataset (when smaller scores indicate better matches).

Match-specific factors include the number of times the reference metabolite of the top match was listed as a candidate match for other query spectra in the dataset (“SpctCndtsCnt”); the number of times the reference metabolite of the top match was identified as the top match for other query spectra in the dataset (“MatchCnt”); and the average cosine similarity score between the spectrum of the matched reference metabolite and all other spectra within the reference library (“RefAvg”). All considered variables were centered and scaled prior to inclusion in models. All possible subsets of these seven factors resulted in 127 models fitted for each of the three GMM extensions, resulting in a total of 381 models compared against the standard GMM model. Models were fitted using the gamlss.mx package (19) in the open-source software environment, R (20).

### 2.4 Similarity Metrics

Several similarity (or equivalently, “distance”) metrics have been developed (21), and many of these metrics have been applied towards GC/MS metabolite identification. However, given this abundance of metrics, there is no current consensus regarding which metric is best, and so it is important that any proposed method for FDR estimation is robust to the choice of score. With this in mind, we therefore consider 28 different similarity metrics in the evaluations of our proposed FDR estimation methodology. These scores include the cosine correlation, which is the normalized inner product of the two vectors representative of compared spectra (21); the Stein-Scott similarity metric that incorporates peak intensity and other MS-specific information in its computation (22); and the squared Euclidean distance, which is computed as the sum of squared differences between the intensities measured at the shared points of two spectra (21). For a full list of the 28 spectra considered, we refer the reader to Table S1 in the supplement.

### 2.5 Comparison Approach

Comparisons between all considered models were made across 812 identification lists generated from the 28 different similarity metrics applied separately to each of the 29 datasets shown in Table 1. Identification lists were generated by selecting the best matching candidate spectra for each query spectrum, according to the given choice of score. These identification lists were then filtered to include only those matches with either a “True Positive” (TP) or “True Negative” (TN) annotation. Given a similarity score threshold *t*, the true FDR was then computed as the proportion of TN annotations among matches with score ≥ *t* (when larger scores are indicative of better matches) or among matches with score ≤ *t* (when smaller scores are indicative of better matches). A true FDR curve, composed of true FDR computations at different values of *t*, was then constructed by using the set of observed scores in the identification lists as the values of *t* (Figure S2). FDR estimates from each model were then obtained at each of the same values of *t* as for the true FDR computations, thus generating estimated FDR curves for each model. To assess the accuracy of each model’s estimates, we computed the absolute value of the difference between each model’s estimated FDR and the true FDR at each value of *t*. For each model, the median of its absolute errors (i.e. MAE) was then used to summarize its FDR estimation accuracy for the given identification list (Figure S3). Distributions of these computed MAEs across all hit-lists were then compared and used to identify the top-performing models relative to baseline. We additionally investigated the use of AIC (23) and BIC (24), popularly used model selection criteria, towards identifying top-performing models but found there to be poor correspondence between AIC-/BIC-preferred models and those that consistently outperform the baseline.

### 2.4 Library Size Simulation

To further strengthen the generalizability of our study results, we designed a simulation study to assess the impact that reference library size has on the relative differences in measured estimation accuracy between the top-performing models identified through the study described in Section 2.3 and the baseline GMM. We simulated libraries that were 25%, 50%, and 75% the size of our original reference library by randomly sampling – without replacement – from our original set of 1284 reference compounds. For each combination of the 29 datasets and 28 similarity scores, 30 sets of libraries of each size were simulated. Candidate lists of potential matches for each spectrum were then filtered to include only the reference metabolites within a particular simulated set, and identification lists were subsequently obtained by identifying the top match among remaining candidates. Post-identification-list-generation, the analyses proceeded in an identical manner as for the comparison study: true and estimated FDR curves were determined, and MAEs were computed as the metric of comparison between models. Code to replicate these and the other analyses described in previous sections is available at All code for plots and analyses are available on Github at https://github.com/PNNL-m-q/metabolomics_fdr.

## 3. RESULTS

### 3.1 Truth Annotation

Table 2 summarizes, by sample type, the average percentage of “True Positive” (TP), “True Negative” (TN), and “Unknown” annotations among the 812 identification lists generated for all analyses. Note that the number of identification lists (i.e ID Lists) for each sample type is computed as the number of datasets for the sample type (Table 1) multiplied by 28, the number of different similarity scores considered.

**Table 2.** Average Annotation Percentages for Identification Lists, Stratified by Sample Type.

| Sample Type | # ID Lists | % TP | % TN | % Unknown |
| --- | --- | --- | --- | --- |
| Standards | 308 | 2.63 | 86.16 | 11.21 |
| Human CSF | 224 | 19.53 | 25.43 | 55.04 |
| Human Blood Plasma | 84 | 15.94 | 14.90 | 69.16 |
| Human Urine | 56 | 7.45 | 19.43 | 73.12 |
| Fungi | 84 | 9.44 | 31.05 | 59.51 |
| Soil | 56 | 9.83 | 33.13 | 57.04 |

We observe (on average) that for all but the standards samples, more than half of all identifications within generated ID lists are labeled as “Unknown” and thus removed from analyses. Based on these average percentages, the number of spectra informing the results of subsequent analyses are (roughly) 9618 standards spectra; 11265 human CSF spectra; 4842 human blood plasma spectra; 6204 human urine spectra; 3506 fungi spectra; and 1557 soil spectra.

### 3.2 Model Comparison

Recall that the comparisons between the 382 considered models are based on the median absolute estimation error (MAE) of estimated FDR curves relative to the truth. These MAEs are computed for each of the 812 ID lists, and thus each model is associated with 812 different MAEs. Subsetting by sample type and faceting by candidate model extension type, each model’s median observed MAE is then represented within the distributions shown in Figure 2.

**Figure 2.**
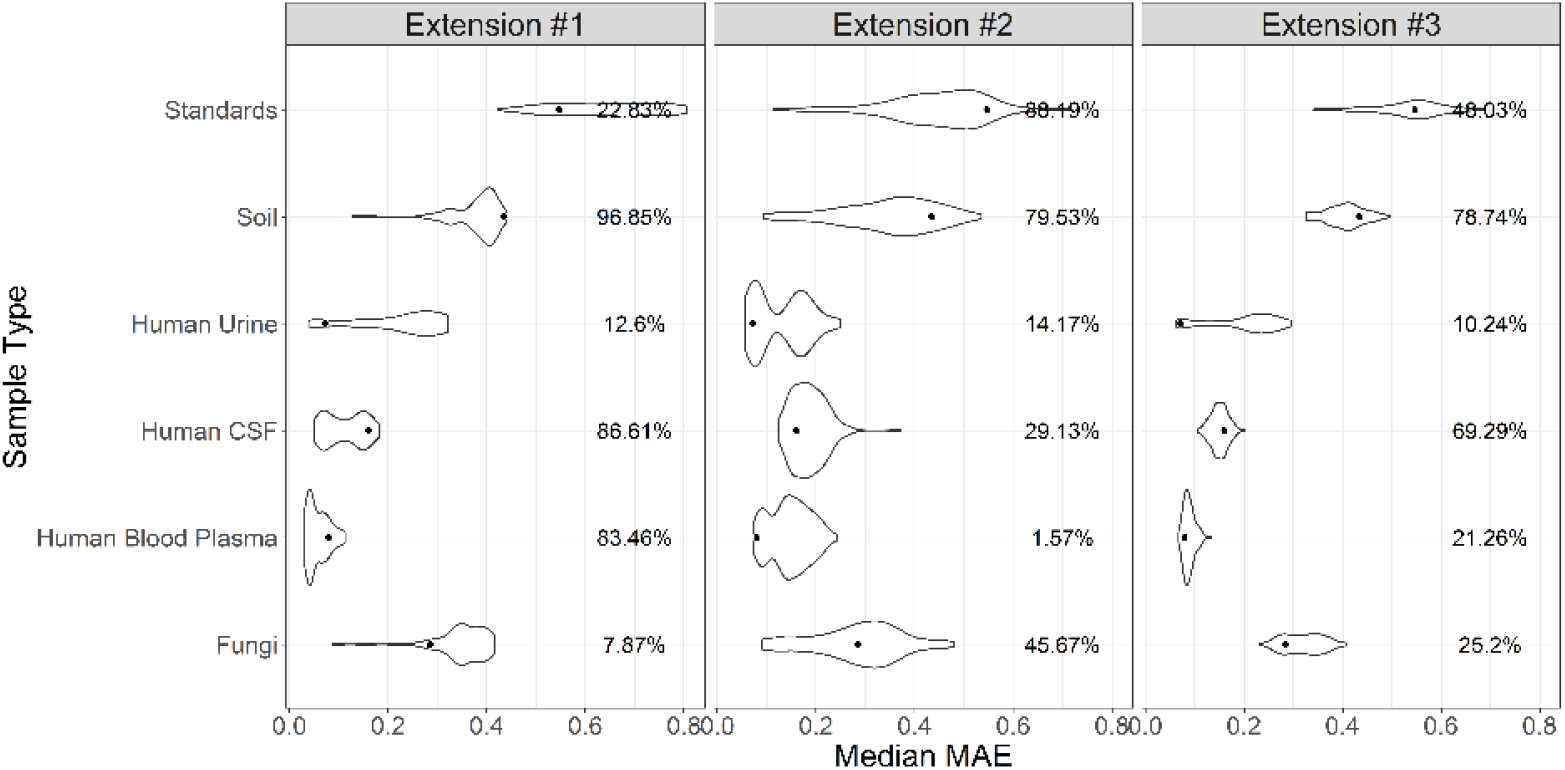
Distributions of model median MAE values by sample type, faceted by extension type. Each distribution is formed by the median MAEs recorded for each of the 382 compared models (127 for each extension). Black points are representative of the standard GMM median MAE value under the corresponding sample type. The percentages of models with a smaller (improved) median MAE relative to baseline are provided to the right of each plotted distribution. Note that a median MAE of 1 implies an FDR estimation error of 100%.

The displayed distributions indicate that for each sample type, there are candidate models from each extension whose performance is improved relative to baseline. However, the number of improved candidate models varies by extension and sample type. As an example, for standards samples, we observe that 88.19% of the “Extension #2” candidate models have smaller median MAEs than the standard GMM, yet only 22.83% of the “Extension #1” models are improved over baseline. This trend is reversed for human blood plasma samples, where 83.46% of “Extension #1” models and only 1.57% of “Extension #2” models are improved. The degree of relative improvement among candidate models, measured by the difference between each distribution’s minimum median MAE and that of the baseline GMM, also exhibits substantial variability across sample type. For example, in soil samples, these differences are roughly 0.32 (“Extension #1), 0.35 (“Extension #2”), and 0.125 (“Extension #3”), contrasting with the human urine samples differences of ~0.03 (“Extension #1”), ~0.02 (“Extension #2”), and <0.01 (“Extension #3”).

Despite the observed variability of results, Figure 2 nonetheless indicates that there exists subset(s) of candidate models that improve over the baseline GMM model. Our interest, however, is to identify the subset of models whose relative improvements are robust to changes in the underlying sample type and may therefore be applied more generally and reliably as improvements over the standard GMM. To arrive at this subset, we consider several different MAE-based model rankings and identify the model(s) that are consistently among the highest-ranked across the most sample types. Ranked lists of candidate models were generated based on the 1) average MAE by sample type; 2) median MAE by sample type; 3) a composite rank based on the sum of the average and median MAE ranks; and sample-type-agnostic ranks based on the 4) average and 5) median MAE. Only the top 4 (~1%) ranked models were recorded for each of the five ranking methods considered. Across all resulting top-ranked subsets, three candidate models were the most consistently observed. Of these, two were found to be among the top 1% of models for three sample types, and one (“Ext # 1: SpctCndtsCount + ScoreDiff”) was identified as the highest ranked model for two sample types. Figure 3 provides the distributions of the median absolute differences (MAE) between model-estimated and true FDR curves for these three “best” models and the baseline GMM.

**Figure 3.**
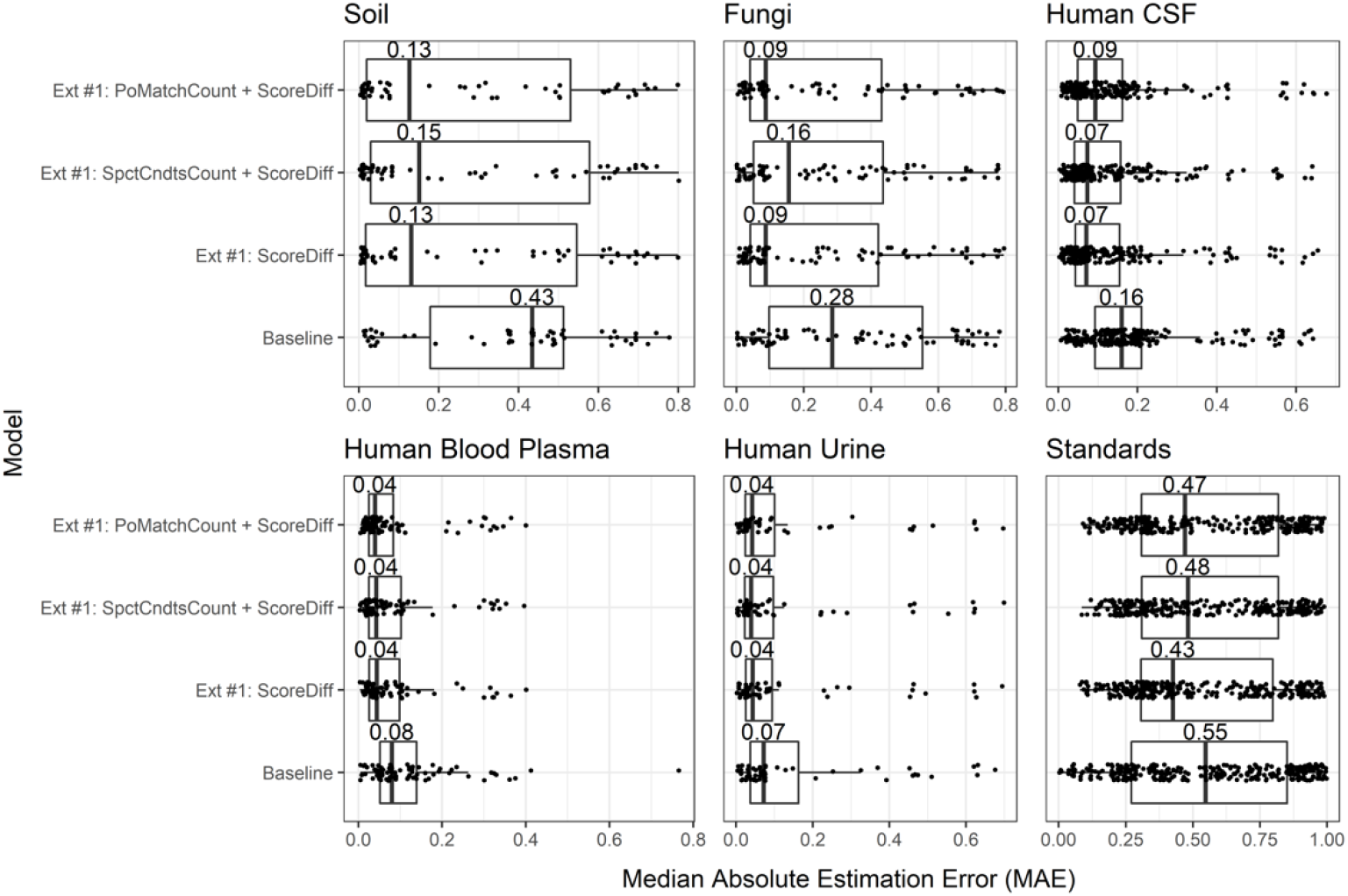
Boxplots of model-specific distributions of the median absolute errors (MAE), faceted by sample type. For a given boxplot, black points indicate the MAEs of the underlying samples (i.e. identification lists) that generate the distribution. Each boxplot is annotated by its median value (i.e. the median MAE).

As was the case for the comparisons depicted in Figure 2, we observe here variability across sample types in the improvements that each “best” model exhibits over the baseline GMM. Comparing the median MAEs of the “best” and baseline model distributions, we observe (at worst) a (0.43 – 0.l5)/0.43 * 100 ≈ 65% relative reduction in estimation error for soil samples; 43% for fungi and human urine samples; 44% for human CSF samples; 50% for human blood plasma samples; and 13% for standards samples. Evidence for improvement over baseline is strengthened by the fact that in all but the standards samples, p-values obtained from one-sided Wilcoxon rank-sum tests between the “best” model distributions and the baseline distribution were each less than 0.05. This suggests that – for non-standards samples – the top three models have MAEs that are typically distributed over errors that are smaller than those of the baseline GMM. For standards samples comparisons, two-sample Wilcoxon rank-sum tests for equality indicated an equivalence of MAE distributions and thus “best” models are characterized by errors comparable to the standard GMM in these cases.

### 3.3 Library Size Simulation

Having identified a set of “best” models with relative improvements over the standard GMM that are robust to the choice of sample type and similarity score, Figure 4 next summarizes comparisons between these top models and the baseline across simulated reference libraries of varying size.

**Figure 4.**
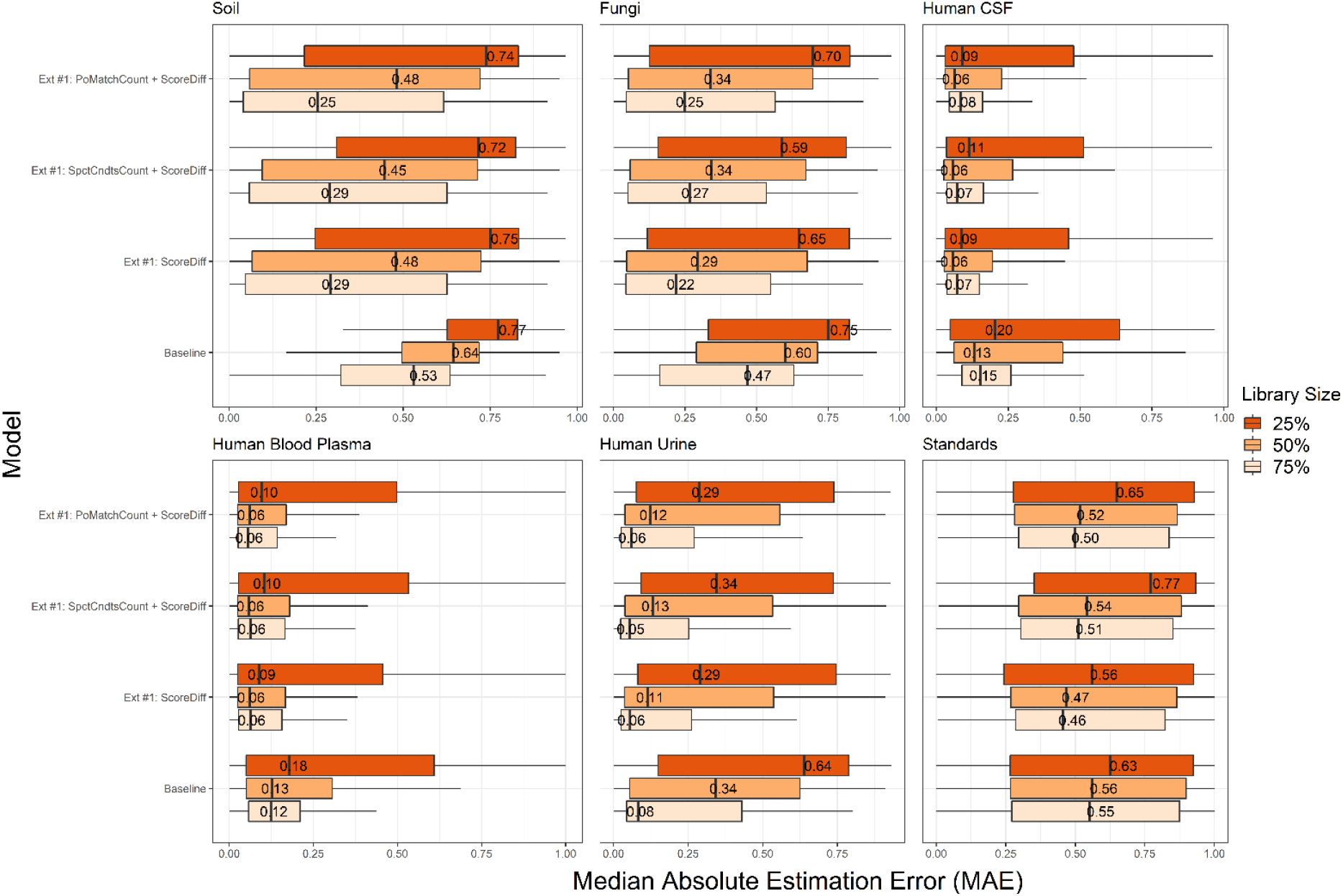
Boxplots of model-specific MAE distributions under simulated reference libraries of varying size, faceted by sample type. Each boxplot is annotated by its median value (i.e. the median MAE). Simulated libraries are either 25%, 50%, or 75% the size of our original reference library of 1284 compounds.

Across the “best” and baseline models, Figure 3 indicates an increase in estimation accuracy with increased library size. This general trend is unaffected by sample type, however the model-specific trends in estimation error improvement do vary across sample type. Taking the baseline model as an example and comparing the median MAEs at simulated libraries that are 25% and 75% of the original library size, we see relative reductions of ~31% for soils samples; ~37% for fungi samples; ~25% for human CSF samples; ~33% for human blood plasma samples; ~88% for human urine samples; and 13% for standards samples.

Comparing across models and at each simulated library size, we see that the previously observed relative improvements (Figure 2) among the “best” models are largely preserved. Furthermore, these “best” models typically experience greater reductions in estimation error with increased library size than what is seen for the baseline model, except for cases where the estimation error is minimal to begin with. Specifically, we observe an average reduction among the “best” models of ~62% for soils and fungi samples (baseline GMM: 31% and 37%); ~23% for human CSF samples (baseline GMM: 25%); ~38% for human blood plasma (baseline GMM: 33%); ~81% for human urine samples (baseline GMM: 88%); and 25% for standards samples (baseline GMM: 13%) when increasing the simulated library size from 25% to 75% of the size of our original reference library.

## 4. DISCUSSION

In aggregate, the results from our studies provide support towards three models as improved alternatives over the standard GMM for FDR estimation within GC/MS metabolomics. These three models are each of the first extension type, which allows the mixing proportions of the latent true and false positive populations to vary by some set of covariates. More practically speaking, models of the first extension type are those that assume that there are a greater (or lesser) proportion of true positives within a set of identifications, depending on some set of external factor(s). The factors characterizing the three identified top models include “PoMatchCount”, “ScoreDiff”, and “SpctCndtsCount”, with “ScoreDiff’ being common to all three top models.

The consistent presence of “ScoreDiff’ across all three of our top-performing models agrees with the findings of Kim and Zhang (9) regarding similarity score differences and FDR estimation. The idea that “ScoreDiff’ offers value towards FDR estimation also agrees with intuition: smaller differences between the top two scoring candidate matches imply the existence of at least one other supported match for the same spectrum, thus creating greater uncertainty for the match. Given the importance that score differences exhibit towards FDR estimation, future work may investigate the utility of using score differences – rather than the scores of the top matches – as the basis for standard GMM models. This proposed work may lead to yet another simple and improved alternative to the standard GMM, which has the potential to be improved even further by leveraging the GMM extensions proposed in the present work.

Like “ScoreDiff”, the importance of each of the other observed factors in our top models makes intuitive sense. To understand this, it is important to emphasize that each of these factors are used to model the mixing proportion between the latent true and false positive populations. In other words, depending on the values of these factors, the model-estimated proportion of true positives may be higher or lower. Considering “PoMatchCount”, which describes the number of candidate matches returned for a given spectrum, larger values indicate a greater number of potential matches and thus a larger proportion of false positives (assuming only a single match is true). The metabolite-specific factor, “SpctCndtsCount”, measures the number of times a reference metabolite appeared as a candidate match across other spectra. This factor may then be thought of as a proxy measure for the ubiquity of a spectrum’s features across known metabolites. If a reference candidate is a potential match across several other spectra, it may have features which are commonly observed and thus its predisposition to be incorrectly matched is higher than a metabolite with more unique features.

The present work makes clear that incorporating information beyond the similarity scores of top matches greatly improves the quality of FDR estimates. Though our top models are characterized by a smaller subset of considered factors, other factors may still hold importance. Recall that our top models are those that were among the best for multiple sample types, not all (or even the majority). This point is best made when comparing the performance of the top models for standards samples in Figure 3 to the best case observed for standards samples in Figure 2. The lowest median MAE for our top models is 0.43 for standards samples (Figure 3), yet the minimum median MAE observed for standards samples is near 0.10 (Figure 2). The model corresponding to this minimum contains “Interference”, “MatchCount”, “RefAvg”, in addition to all the factors observed in our top models. It should also be noted that this model is of the second GMM extension type, as opposed to the first GMM extension type that characterizes our top models.

Outside of supporting the notion that covariates outside the set identified for our top models may hold importance towards FDR estimation, the previous observation also suggests a potential avenue for future explorations. Ensemble modeling approaches are those that generate estimates or predictions based on an aggregation of several different models. Through leveraging the collective strengths of several models, ensemble approaches have often been found to outperform their non-ensemble counterparts (25). In the present context, given that no single model has been identified as optimal across all sample types, datasets, score choices, and library sizes, an ensemble approach may be ideal. This potential avenue for improvement aside, it should be noted that our present results still demonstrate a robustness across these factors, as many of our proposed models consistently outperformed the baseline comparator under a variety of conditions.

Last, our library size investigations reveal a strong and consistent trend: larger libraries are associated with reduced estimation error. This finding implies that for reference libraries much larger than the library of 1284 compounds used in this study, such as the NIST library, estimation errors may be minimized to an even greater extent than observed here (and vice versa). However, given that the presented library size trends are based on simulated reference libraries, additional work is needed to validate these findings through comparisons based on real reference libraries of varying size.

## 5. CONCLUSION

This work has identified three models that offer improved accuracy over a Gaussian mixture modeling approach when estimating the false discovery rate. Each of these models are easily implemented, as they may be fitted using the well-documented gamlssMXfits function offered by the open-source R package, gamlss.mx. Contrasting the standard GMM, these models estimate the true proportion of correct identifications (i.e. true positives) as a parameter that varies according to a set of external factors. These factors include the number of candidate matches returned for the given spectrum, the number of times a matched metabolite appeared as a candidate match for other spectra, and the difference in similarity score between the top two candidate matches for the given spectrum. The relative improvements in performance exhibited by these models have been shown to be robust to the choice of sample type, dataset, similarity score, and library size. However, there has been a clear contrast in performance between standards and non-standards samples. Whereas estimation error for the top three identified models is on the order of ~10% or lower for non-standards samples, typical estimation errors rise fourfold for standards samples. A similar inflation in estimation error is observed more generally across all models. This observation highlights the importance of using real, non-standards data when developing methodology for false discovery rate estimation. Standards samples do not reflect the characteristics and complexities of real-world biological samples and thus methods that may be optimal for standards data may be sub-optimal otherwise.

## ASSOCIATED CONTENT

### Supporting Information

GMM/HEBM simulation study description and key finding; supplemental figure depicting the true FDR curves of all identification lists; supplemental figure describing computation of the median absolute estimation error. A supplemental table providing information of similarity metrics used in model comparisons. This material is available free of charge via the Internet at http://pubs.acs.org.

## AUTHOR INFORMATION

### Author Contributions

VLP prepared standards and acquired some of the data used; YEC developed the software used for the metabolite identification and provided the data; DJD, LMB, and JEF developed the statistical analysis pipeline with input from all authors; CSC provided the truth annotations and supervised the project; JEF wrote the manuscript with significant contributions from all authors

## ACKNOWLEDGMENT

A portion of this research was performed using data acquired by the Environmental Molecular Sciences Laboratory, a DOE Office of Science User Facility sponsored by the Biological and Environmental Research program under Contract No. DE-AC05-76RL01830. The authors would like to thank Allison Thompson for contribution towards the programming of similarity scoring metrics in CoreMS.

## ABBREVIATIONS

GC-MS: gas chromatography - mass spectrometry;
FDR: false discovery rate;
GMM: Gaussian mixture model;
MAE: median absolute estimation error;
LC-MS/MS: liquid chromatography - tandem mass spectrometry;
NIST: National Institute of Standards and Technology;
PNNL: Pacific Northwest National Laboratory;
TMS: trimethylsilyation;
FAME: fatty acid methyl ester;
CSF: cerebrospinal fluid;
TP: true positive;
TN: true negative;
RI: retention index.

## REFERENCES

1. Matsuda, F.; Tsugawa, H.; Fukusaki, E. Method for Assessing the Statistical Significance of Mass Spectral Similarities Using Basic Local Alignment Search Tool Statistics. Anal Chem 2013, 85 (17), 8291–8297.

2. Elias, J.; Gygi, S. Target-decoy search strategy for increased confidence in large-scale protein identifications by mass spectrometry. Nat Methods 2007, 4, 207–214. https://doi.org/10.1038/nmeth1019

3. Scheubert, K.; Hufsky, F.; Petras, D.; Wang, M.; Nothias, L.; Dührkop, K.; Bandeira, N.; Dorrestein, P. C.; Böcker, S. Significance estimation for large scale metabolomics annotations by spectral matching. Nat Commun 2017, 8, 1494. https://doi.org/10.1038/s41467-017-01318-5

4. Rasche, F.; Svatoš, A.; Maddula, R. K.; Böttcher, C.; Böcker, S. Computing fragmentation trees from tandem mass spectrometry data. Anal. Chem. 2011, 83, 1243–1251.

5. Böcker, S.; Dührkop, K. Fragmentation trees reloaded. J Cheminform 2016, 8, 5. https://doi.org/10.1186/s13321-016-0116-8

6. Efron, B.; Tibshirani, R.; Storey, J. D.; Tusher, V. Empirical Bayes analysis of a microarray experiment. J. Am. Stat. Assoc 2001, 96, 1151–1160.

7. Stein S. E. Estimating probabilities of correct identification from results of mass spectral library searches. Journal of the American Society for Mass Spectrometry, 1994, 5(4), 316–323. https://doi.org/10.1016/1044-0305(94)85022-4

8. Jeong, J.; Shi, X.; Zhang, X.; Kim, S.; Shen, C. An empirical Bayes model using a competition score for metabolite identification in gas chromatography mass spectrometry. BMC Bioinformatics 2011, 12.

9. Kim, S.; Zhang, X. Discovery of false identification using similarity difference in GC-MS-based metabolomics. J Chemometr 2015, 29 (2), 80–86.

10. Corilo, Y. E.; Kew, W. R.; McCue, L. (2021, March 27). EMSL-Computing/CoreMS: CoreMS 1.0.0 (Version v1.0.0), as developed on Github. Zenodo. http://doi.org/10.5281/zenodo.4641553

11. Kind, T.; Wohlgemuth, G.; Lee, D. Y.; Lu, Y.; Palazoglu, M.; Shahbaz, S.; Fiehn, O. FiehnLib: mass spectral and retention index libraries for metabolomics based on quadrupole and time-of-flight gas chromatography/mass spectrometry. Anal Chem 2009, 81 (24), 10038–48.

12. Sharon, G.; Cruz, N. J.; Kang, D.; Gandal, M. J.; Wang, B.; Kim, Y.; Zink, E. M.; Casey, C. P.; Taylor, B. C.; Lane, C. J.; Bramer, L. M.; Isern, N. G.; Hoyt, D. W.; Noecker, C.; Sweredoski, M. J.; Moradian, A.; Borenstein, E.; Jansson, J. K.; Knight, R.; Metz, T. O.; Lois, C.; Geschwind, D. H.; Krajmalnik-Brown, R.; Mazmanian, S. K. Human Gut Microbiota from Autism Spectrum Disorder Promote Behavioral Symptoms in Mice. Cell 2019, 177(6), 1600–1618.e17, https://doi.org/10.1016/j.cell.2019.05.004

13. Ulrich, D.E.M.; Clendinen, C.S.; Alongi, F. et al. Root exudate composition reflects drought severity gradient in blue grama (Bouteloua gracilis). Sci Rep 2022, 12, 12581. https://doi.org/10.1038/s41598-022-16408-8

14. Kyle, J.E.; Stratton, K.G.; Zink, E.M. et al. A resource of lipidomics and metabolomics data from individuals with undiagnosed diseases. Sci Data 2021, 8(114), https://doi.org/10.1038/s41597-021-00894-y

15. Wishart, D. S.; Guo, A.; Oler, E.; Wang, F.; Anjum, A.; Peters, H.; Dizon, R.; Sayeeda, Z.; Tian, S.; Lee, B. L.; Berjanskii, M.; Mah, R.; Yamamoto, M.; Jovel, J.; Torres-Calzada, C.; Hiebert-Giesbrecht, M.; Lui, V. W.; Varshavi, D.; Varshavi, D.; Allen, D.; … Gautam, V. HMDB 5.0: the Human Metabolome Database for 2022. Nucleic acids research 2022, 50(D1), D622–D631. https://doi.org/10.1093/nar/gkab1062

16. Gil de la Fuente, A.; Godzien, J.; Fernández López, M.; Rupérez, F. J.; Barbas, C.; Otero, A. Knowledge-based metabolite annotation tool: CEU Mass Mediator. Journal of Pharmaceutical and Biomedical Analysis 2018, 154, 138–149. https://doi.org/10.1016/j.jpba.2018.02.046

17. Dempster, A. P.; Laird, N. M.; Rubin, D. B. Maximum Likelihood from Incomplete Data via the EM Algorithm. Journal of the Royal Statistical Society. Series B (Methodological) 1977, 39 (1), 1–38. http://www.jstor.org/stable/2984875

18. Rigby, R. A.; Stasinopoulos, D. M. Generalized Additive Models for Location, Scale and Shape. Journal of the Royal Statistical Society: Series C (Applied Statistics) 2005, 54 (3), 507–554.

19. Stasinopoulos, M.; Rigby B. (2020). gamlss.mx: Fitting Mixture Distributions with GAMLSS. R package version 6.0-0. https://CRAN.R-project.org/package=gamlss.mx

20. R Core Team (2021). R: A language and environment for statistical computing. R Foundation for Statistical Computing, Vienna, Austria. URL https://www.R-project.org/

21. Cha, S. Comprehensive Survey on Distance/Similarity Measures between Probability Density Functions. International Journal of Mathematical Models and Methods in Applied Sciences, 2006, 1. https://tcs.ah-epos.eu/eprints/1372/

22. Stein, S. E., & Scott, D. R. Optimization and testing of mass spectral library search algorithms for compound identification. Journal of the American Society for Mass Spectrometry, 1994, 5(9), 859–866. https://doi.org/10.1016/1044-0305(94)87009-8

23. Akaike, H. A new look at the statistical model identification. IEEE Transactions on Automatic Control 1974, 19 (6), 716–723, doi: 10.1109/TAC.1974.1100705

24. Schwarz, G. Estimating the Dimension of a Model. The Annals of Statistics 1978, 6(2), 461–464. http://www.jstor.org/stable/2958889

25. Sagi, O; Rokach, L. Ensemble learning: A survey. WIREs Data Mining Knowl Discov. 2018, 8:e1249. https://doi.org/10.1002/widm.1249

