## Supplemental Data 1 for "Gaussian Mixture Modeling Extensions for Improved False Discovery Rate Estimation in GC-MS Metabolomics"

### SUPPLEMENTAL MATERIALS

**S1. GMM Simulation Study Description.** We compared the false discovery rate estimation accuracy of the standard Gaussian Mixture model (GMM) to that of the hierarchical empirical Bayes model (HEBM) introduced by Jeong et. al (8). Data for this comparison study were simulated according to the HEBM. To simulate data from the HEBM, we first specified values for each of the model's parameters ($\rho = 0.1, \eta_0 = 0.5, \beta_0 = 3, \beta_1 = 2, \beta_2 = 1, \eta_1 = 0.7, \alpha_0 = 6, \alpha_1 = 4, \alpha_2 = 2, \tau = 0.8, \mu_T = 5, \sigma_T^2 = 9, \mu_F = 40, \sigma_F^2 = 100$). Next, we computed competition scores based on a subset of 200 compounds from the CoreMS reference library. We then drew 200 samples from a $Bernoulli(\rho)$ distribution to generate a binary vector $Y$ indicating which of the 200 reference metabolites are "present" ($Y_i = 1$) in the simulated sample. Using the generated vector, $Y$, we then take draws from two different Bernoulli distributions, depending on whether $Y_i = 1$ or $Y_i = 0$. The parameterizations of these Bernoulli distributions are functions of the computed competition scores, and the parameters $\eta_0, \beta_0, \beta_1, \beta_2, \eta_1, \alpha_0, \alpha_1$, and $\alpha_2$. Drawing from this second set of Bernoulli distributions results in the binary vector $Z$ indicating which of the 200 reference metabolites were "matched" ($Z_i = 1$) to simulated sample spectra. Each "matched" reference metabolite (i.e. $Z_i = 1$) is then associated with five hypothetical sample spectra. For each of these five hypothetical matches, we next draw from a $Bernoulli(\tau)$ distribution if the corresponding reference metabolite is "present" within the simulated sample ($Y_i = 1$). If not present, we draw from a $Bernoulli(0)$ distribution. This process results in the generation of binary vectors $W_i$ that indicate whether the "match" between the $i^{th}$ reference metabolite and $j^{th}$ hypothetical match is correct ($W_{ij} = 1$). Last, the cosine similarity scores between the reference metabolite and its simulated matches are generated from a normal mixture distribution with parameterization depending on the value of $W_{ij}$. If $W_{ij} = 0$, scores are generated from the "false positive" component of the mixture distribution parameterized by $\mu_F$ and $\sigma_F^2$. Otherwise, scores are generated from the "true positive" component of the mixture distribution parameterized by $\mu_T$ and $\sigma_T^2$. Further details on the HEBM, its structure, required parameters, and the competition scores may be found in Jeong et. al (8).

A total of 100 datasets were simulated based on the previously described process, and for each, GMM and HEBM models were fitted to the data to yield model-estimated FDR curves. The HEBM outputs probabilities indicating the likelihood of each reference metabolite's presence in sample, $\Pr(Y_i = 1)$, and thus FDR estimates for the HEBM are based on thresholding these probabilities (e.g. all reference metabolites with estimated probabilities greater than 0.8 are considered "identifications"). This differs from the process of obtaining FDR estimates based on the GMM. Details on FDR estimation for the GMM may be found in the main text. True FDR curves for each dataset were obtained by determining the proportion of false positives ($Y_i = 0$) among identifications generated according to each observed score threshold (GMM) or probability threshold (HEBM). Figure S1 displays the resulting model-estimated and true FDR curves for each of the 100 simulated datasets.

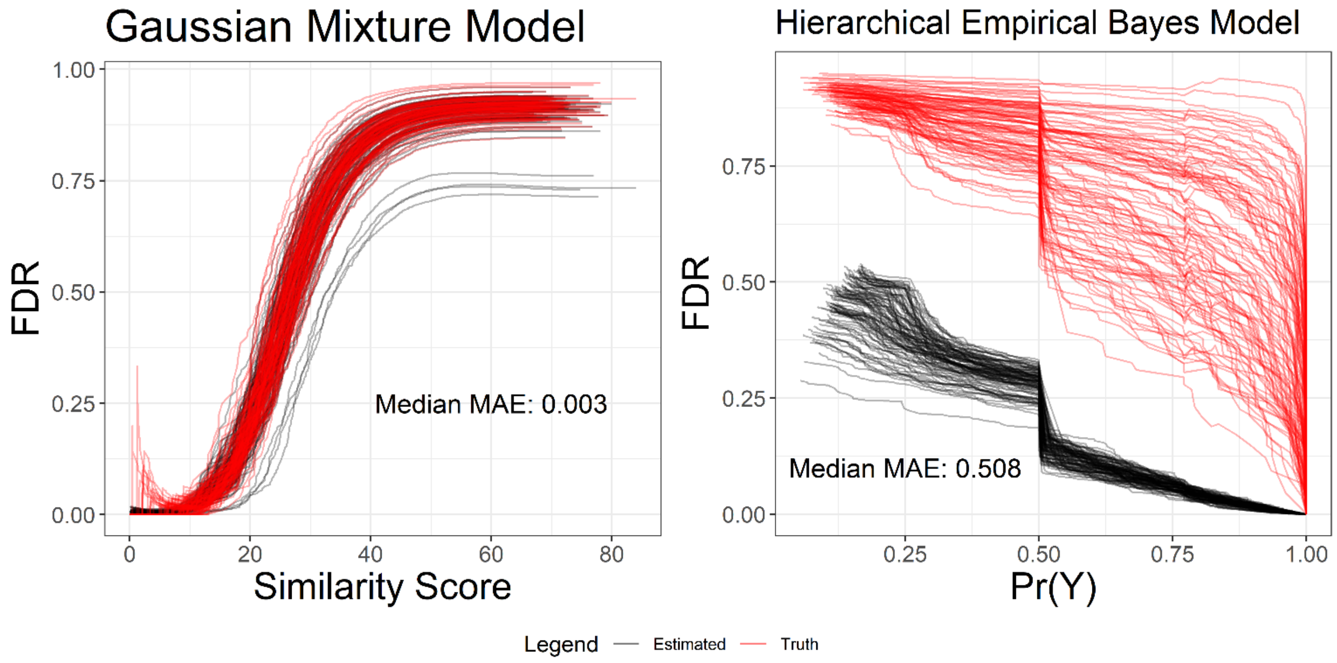


**Figure S1**. Model-estimated FDR curves based on the Gaussian mixture model (left) or hierarchical empirical Bayes model (right). Each black line corresponds to the model-estimated curve of a simulated dataset. Red lines indicate the true FDR curves of each dataset. The text indicates the median of the median absolute errors (MAE) measured between each estimated FDR curve and the truth across the 100 simulated datasets.

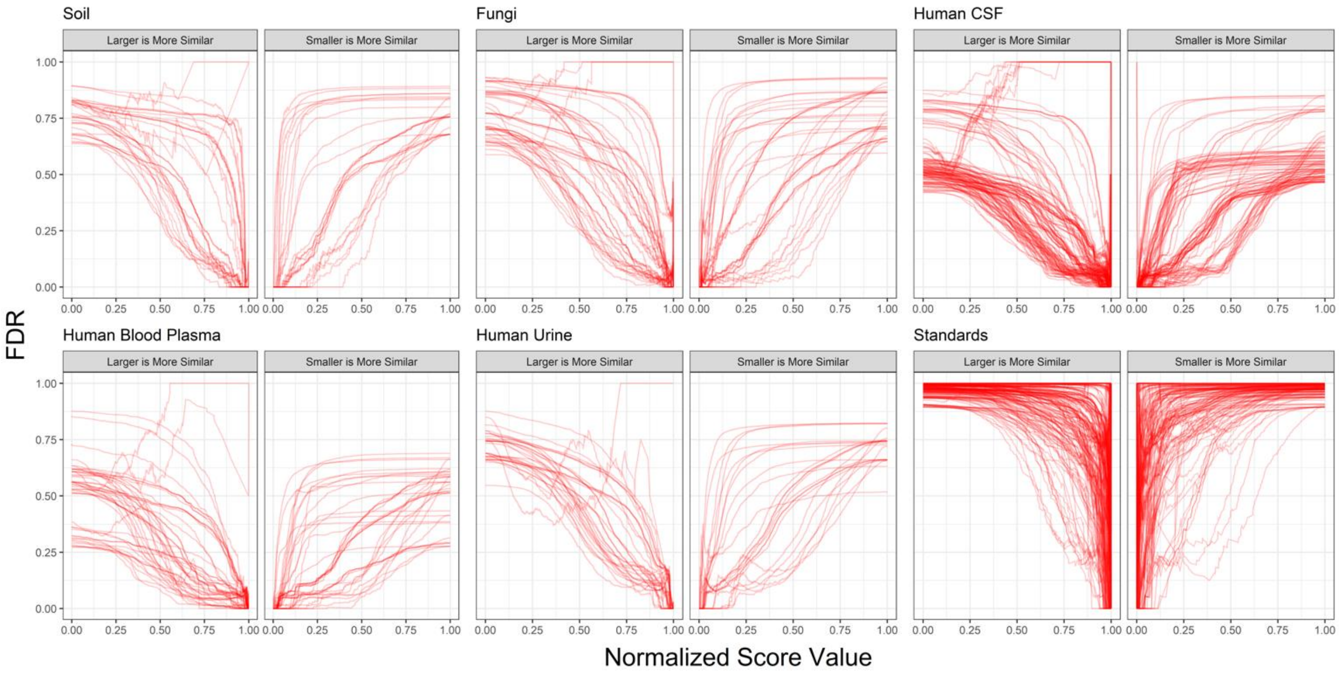


**Figure S2**. True false discovery rate (FDR) curves corresponding to each of the 812 identification lists, stratified by sample type. Recall that the 812 identification lists result from applying each of the 28 similarity metrics to each of the 29 datasets. Thus, each curve represents a particular metric/dataset combination. Similarity metrics are separated according to whether larger or smaller values indicate improved similarity (as indicated by the subplot facets). To plot all curves on the same scale, all similarity metrics have been normalized to fall within the range of 0 and 1.

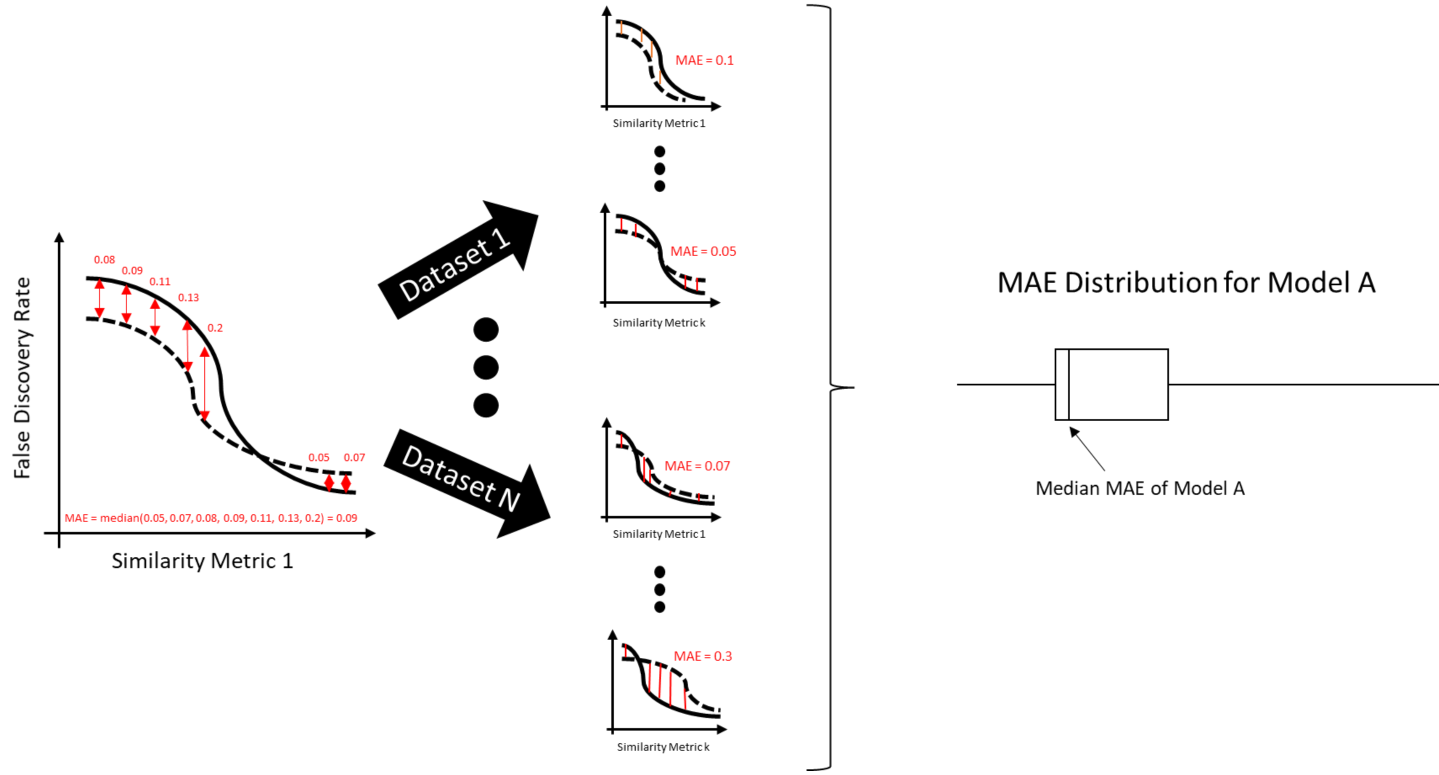


**Figure S3.** Visual description of measurements used to assess the accuracy of model-generated FDR estimates. For each of N datasets, k similarity metrics are considered to generate sets of true (solid line) and model-estimated (dashed line) FDR curves corresponding to the particular dataset/similarity-score combination. Only a single estimated curve is shown in the figure above, but estimated curves are obtained for every considered model. The median of the absolute value of differences between estimated and true curves (MAE) along observed threshold points is used to summarize the estimation accuracy of a model for a particular dataset/similarity-score combination. Collectively, measured MAEs form a distribution which may be used to describe a model's overall performance across different choices of dataset and similarity metric.

**Table S1. Similarity and Distance Metrics Used in Model Comparisons**

| Metric | Directionality | Reference |
|---|---|---|
| Similarity Score | Larger scores indicate better matches | Koo I, Kim S, Zhang X (2013). Comparative analysis of mass spectral matching-based compound identification in gas chromatography-mass spectrometry. J Chromatogr A. 1298:132-8. doi: 10.1016/j.chroma.2013.05.021. Epub 2013 May 13. PMID: 23726352; PMCID: PMC3686837. |
| Spectral Similarity Score | Larger scores indicate better matches | Cha, S. (2006). Comprehensive Survey on Distance/Similarity Measures between Probability Density Functions. International Journal of Mathematical Models and Methods in Applied Sciences, 1. https://tcs.ah-epos.eu/eprints/1372/ |
| Weighted Cosine Correlation | Larger scores indicate better matches | Koo I, Kim S, Zhang X (2013). Comparative analysis of mass spectral matching-based compound identification in |

| | | |
|---|---|---|
| | | gas chromatography-mass spectrometry. J Chromatogr A. 1298:132-8. doi: 10.1016/j.chroma.2013.05.021. Epub 2013 May 13. PMID: 23726352; PMCID: PMC3686837. |
| Cosine Correlation | Larger scores indicate better matches | Cha, S. (2006). Comprehensive Survey on Distance/Similarity Measures between Probability Density Functions. International Journal of Mathematical Models and Methods in Applied Sciences, 1. https://tcs.ah-epos.eu/eprints/1372/ |
| Stein Scott Similarity | Larger scores indicate better matches | Stein, S. E., & Scott, D. R. (1994). Optimization and testing of mass spectral library search algorithms for compound identification. Journal of the American Society for Mass Spectrometry, 5(9), 859–866. https://doi.org/10.1016/1044-0305(94)87009-8 |
| Stein Scott Similarity (NIST) | Larger scores indicate better matches | Stein, S. E., & Scott, D. R. (1994). Optimization and testing of mass spectral library search algorithms for compound identification. Journal of the American Society for Mass Spectrometry, 5(9), 859–866. https://doi.org/10.1016/1044-0305(94)87009-8 |
| Pearson Correlation | Larger scores indicate better matches | Pearson, K. (1895). VII. Note on regression and inheritance in the case of two parents. Proceedings of the Royal Society of London, 58(347–352), 240–242. https://doi.org/10.1098/rspl.1895.0041 |
| Spearman Correlation | Larger scores indicate better matches | Fieller, E. C., Hartley, H. O., & Pearson, E. S. (1957). Tests for Rank Correlation Coefficients. I. Biometrika, 44(3–4), 470–481. https://doi.org/10.1093/biomet/44.3-4.470 |
| DWT Correlation | Larger scores indicate better matches | Koo I, Kim S, Zhang X (2013). Comparative analysis of mass spectral matching-based compound identification in gas chromatography-mass spectrometry. J Chromatogr A. 1298:132-8. doi: 10.1016/j.chroma.2013.05.021. Epub 2013 May 13. PMID: 23726352; PMCID: PMC3686837. |
| DFT Correlation | Larger scores indicate better matches | Koo I, Kim S, Zhang X (2013). Comparative analysis of mass spectral matching-based compound identification in gas chromatography-mass spectrometry. J Chromatogr A. 1298:132-8. doi: 10.1016/j.chroma.2013.05.021. Epub 2013 May 13. PMID: 23726352; PMCID: PMC3686837. |
| Chebyshev Distance | Smaller scores indicate better matches | Cha, S. (2006). Comprehensive Survey on Distance/Similarity Measures between Probability Density Functions. |

| | | |
|---|---|---|
| | | International Journal of Mathematical Models and Methods in Applied Sciences, 1. https://tcs.ah-epos.eu/eprints/1372/ |
| Squared Euclidean Distance | Smaller scores indicate better matches | Cha, S. (2006). Comprehensive Survey on Distance/Similarity Measures between Probability Density Functions. International Journal of Mathematical Models and Methods in Applied Sciences, 1. https://tcs.ah-epos.eu/eprints/1372/ |
| Fidelity Similarity | Larger scores indicate better matches | Cha, S. (2006). Comprehensive Survey on Distance/Similarity Measures between Probability Density Functions. International Journal of Mathematical Models and Methods in Applied Sciences, 1. https://tcs.ah-epos.eu/eprints/1372/ |
| Squared-chord Distance | Smaller scores indicate better matches | Cha, S. (2006). Comprehensive Survey on Distance/Similarity Measures between Probability Density Functions. International Journal of Mathematical Models and Methods in Applied Sciences, 1. https://tcs.ah-epos.eu/eprints/1372/ |
| Topsoe Distance | Smaller scores indicate better matches | Cha, S. (2006). Comprehensive Survey on Distance/Similarity Measures between Probability Density Functions. International Journal of Mathematical Models and Methods in Applied Sciences, 1. https://tcs.ah-epos.eu/eprints/1372/ |
| Canberra Metric | Smaller scores indicate better matches | Cha, S. (2006). Comprehensive Survey on Distance/Similarity Measures between Probability Density Functions. International Journal of Mathematical Models and Methods in Applied Sciences, 1. https://tcs.ah-epos.eu/eprints/1372/ |
| Hellinger Distance | Smaller scores indicate better matches | Cha, S. (2006). Comprehensive Survey on Distance/Similarity Measures between Probability Density Functions. International Journal of Mathematical Models and Methods in Applied Sciences, 1. https://tcs.ah-epos.eu/eprints/1372/ |
| Dice Similarity | Larger scores indicate better matches | Cha, S. (2006). Comprehensive Survey on Distance/Similarity Measures between Probability Density Functions. International Journal of Mathematical Models and Methods in Applied Sciences, 1. https://tcs.ah-epos.eu/eprints/1372/ |
| Jensen Differences Distance | Smaller scores indicate better matches | Cha, S. (2006). Comprehensive Survey on Distance/Similarity Measures between Probability Density Functions. International Journal of Mathematical Models and Methods in Applied Sciences, 1. https://tcs.ah-epos.eu/eprints/1372/ |
| Battacharya Distance | Smaller scores indicate better matches | Cha, S. (2006). Comprehensive Survey on Distance/Similarity Measures between |

| | | |
|---|---|---|
| | | Probability Density Functions. International Journal of Mathematical Models and Methods in Applied Sciences, 1. https://tcs.ah-epos.eu/eprints/1372/ |
| Gower Distance | Smaller scores indicate better matches | Cha, S. (2006). Comprehensive Survey on Distance/Similarity Measures between Probability Density Functions. International Journal of Mathematical Models and Methods in Applied Sciences, 1. https://tcs.ah-epos.eu/eprints/1372/ |
| Improved Square Root Cosine Similarity | Larger scores indicate better matches | Sohangir, S., & Wang, D. (2017). Improved sqrt-cosine similarity measurement. Journal of Big Data, 4(1). https://doi.org/10.1186/s40537-017-0083-6 |
| Intersection Similarity | Larger scores indicate better matches | Cha, S. (2006). Comprehensive Survey on Distance/Similarity Measures between Probability Density Functions. International Journal of Mathematical Models and Methods in Applied Sciences, 1. https://tcs.ah-epos.eu/eprints/1372/ |
| Squared Chi-Squared Distance | Smaller scores indicate better matches | Cha, S. (2006). Comprehensive Survey on Distance/Similarity Measures between Probability Density Functions. International Journal of Mathematical Models and Methods in Applied Sciences, 1. https://tcs.ah-epos.eu/eprints/1372/ |
| Squared Root Cosine Correlation | Larger scores indicate better matches | Cha, S. (2006). Comprehensive Survey on Distance/Similarity Measures between Probability Density Functions. International Journal of Mathematical Models and Methods in Applied Sciences, 1. https://tcs.ah-epos.eu/eprints/1372/ |
| Minkowski Distance | Smaller scores indicate better matches | Cha, S. (2006). Comprehensive Survey on Distance/Similarity Measures between Probability Density Functions. International Journal of Mathematical Models and Methods in Applied Sciences, 1. https://tcs.ah-epos.eu/eprints/1372/ |
| Kumar Hassebrook Similarity | Larger scores indicate better matches | Cha, S. (2006). Comprehensive Survey on Distance/Similarity Measures between Probability Density Functions. International Journal of Mathematical Models and Methods in Applied Sciences, 1. https://tcs.ah-epos.eu/eprints/1372/ |
| Soergel Distance | Smaller scores indicate better matches | Cha, S. (2006). Comprehensive Survey on Distance/Similarity Measures between Probability Density Functions. International Journal of Mathematical Models and Methods in Applied Sciences, 1. https://tcs.ah-epos.eu/eprints/1372/ |
